# Individual limb measures of split-belt adaptation are correlated with shifts in lower limb position sense

**DOI:** 10.64898/2026.07.19.739475

**Authors:** Jonathan M. Wood, Saunders Penn, Hyosub E. Kim, Susanne M. Morton

**Author notes:** Corresponding author: Jonathan M. Wood, 540 S College Ave, Newark DE, 19713.

## Abstract

Stable gait in humans relies on both precise estimates of lower limb position and the ability to robustly adapt walking movements across different environments. However, the specific relationship between sensorimotor adaptation and the perception of lower limb position during gait remains unclear. In this study, we tested a specific theoretical model of how sensorimotor adaptation during gait could cause a shift in the perception of lower limb position. To test this theory, we probed lower limb position sense estimates using a valid and reliable two-alternative forced choice task before and after a long bout of split-belt treadmill walking. The theoretical model assumes that a shift in the perception of limb position is caused by the optimal integration between the predicted and actual sensory feedback of step lengths. Using this model, we generated hypotheses relating to the direction of the shift in limb position sense estimates as well as the relationship between the magnitude of these sensory shifts and the plateau of adaptation when measured using the canonical step length asymmetry index. Like prior studies, participants robustly adapted their step length asymmetry during split-belt treadmill walking. However, we did not observe reliable changes in limb position sense after this adaptation relative to baseline. Furthermore, the changes in limb position sense were not related to the plateau of sensorimotor adaptation. However, post-hoc analysis of other gait parameters, most notably double limb support time, revealed a strong relationship between its adaptation plateau and sensory recalibration. These results suggest that while the specific theorical model we proposed was not supported, sensory recalibration could play an important role in split-belt adaptation and raise questions regarding the error signal driving split-belt adaptation.

## Introduction

Efficient human locomotion requires navigation over various surfaces, with different footwear, levels of fatigue or pain, and changes in our bodies as we grow. The process that keeps movements calibrated across these circumstances is known as sensorimotor adaptation [1]. For example, when transitioning from walking on a boardwalk to a sandy beach, subtle adjustments to motor commands (i.e., motor recalibration) help maintain stable gait on the more compliant surface. Stable gait also relies on accurate and precise estimates of lower limb position [2–7]. Along with motor recalibration, recalibration of limb position estimates (i.e., sensory recalibration) occurs during sensorimotor adaptation of reaching movements [8–13]. However, despite the importance of limb position sense during walking, no studies have demonstrated evidence for sensory recalibration during gait, or if there is a relationship between sensory recalibration and sensorimotor adaptation itself. Understanding the connection between sensory recalibration and sensorimotor adaptation during gait is critical for refining models of this form of motor learning and could have important clinical implications given the number of individuals with neurologic diagnoses, like stroke, who also have impaired limb position sense.

Sensorimotor adaptation during gait is often studied using the split-belt treadmill (i.e., split-belt adaptation), where two belts, one under each limb, move at different speeds, causing asymmetry of several gait parameters (e.g., step length, double limb support time, limb angle phasing, center of pressure) which are recalibrated to restore symmetric walking [1,14,15]. In addition, perception of lower limb velocity is also recalibrated after split-belt adaptation [16–18]. In these studies, perception of limb velocity was measured during walking by asking participants to manually adjust the speed of one of the two treadmill belts until it matched the perceived speed of the other belt. After split-belt adaptation, participants consistently sensed that the limb that was on the fast belt moved slower compared to its actual speed. However, because this test was always performed during walking, it is unclear the extent to which motor recalibration contributes to the measurement of perceived limb velocity.

Two prior studies have measured lower limb position sense before and after split-belt adaptation to isolate sensory recalibration from motor recalibration as best as possible [18,19]. In these studies, participants judged limb position after the split-belt passively moved either one or both limbs to various locations. Passively moving the limb is thought to prevent motor recalibration from influencing the limb position estimates and reduces the ability of the efference copy (i.e., the predicted sensory consequences of the movement) from influencing position sense estimates [20–22]. While neither study observed sensory recalibration, this could be due to measurement confounds [23], or perhaps the split-belt adaptation period was too short [24]. Of course, it is also possible sensory recalibration does not occur during gait [25]. However, this theory disagrees with findings in sensorimotor adaptation during reaching. For example, in visuomotor adaptation, where a cursor representing the hand position is offset during reaching movements, sensory recalibration is driven by the mismatch between sensory modalities (i.e., vision and proprioception) [9,25–27]. Sensory recalibration also occurs during force field adaptation, where there is no mismatch between visual and proprioceptive feedback [11,12,28–30]. Sensory recalibration in force field adaptation has been hypothesized to be caused by a mismatch between the predicted sensory consequences of the movement and the sensory feedback received from the movement [25,27]. To resolve this mismatch and create a unified percept of limb position, the system could optimally integrate the sensory prediction with the actual sensory feedback of the limb position [27,31–33]. Like force field adaptation, split-belt adaptation does not cause a mismatch between vision and proprioception. Therefore, assuming the processes governing sensorimotor adaptation are similar in reaching and walking [34–36], we hypothesize that the effects on the sensory system should also be similar.

In Figure 1, we provide a schematic for how sensory recalibration could occur during split-belt adaptation, assuming that the primary error signal is the difference between the step lengths (i.e., step length asymmetry) [1,34,35,37] (but see, [15,24]. When the belts are initially split (Figure 1A, top), the limb on the fast belt (i.e., the fast limb) takes a shorter step than the limb on the slow belt (i.e., the slow limb), while the sensory prediction based on the motor command is that the step lengths should be equal. Optimal integration of the sensory prediction and the sensory feedback would cause sensory recalibration of limb position, resulting in the fast limb feeling more anterior and the slow limb feeling more posterior compared to their true positions (Figure 1B, top). As split-belt adaptation continues, the fast and slow step lengths are driven to be more symmetrical by adjusting the motor commands (Figure 1A, bottom). At the same time, the sensory prediction of these motor commands is being recalibrated by some learning rate (the dashed blue function in Figure 1A, bottom). In this framework, adaptation would plateau when the estimated fast and slow limb positions are equal, in other words, the step lengths are perceived as symmetrical (Figure 1A&B, bottom) [27].

**Figure 1.**
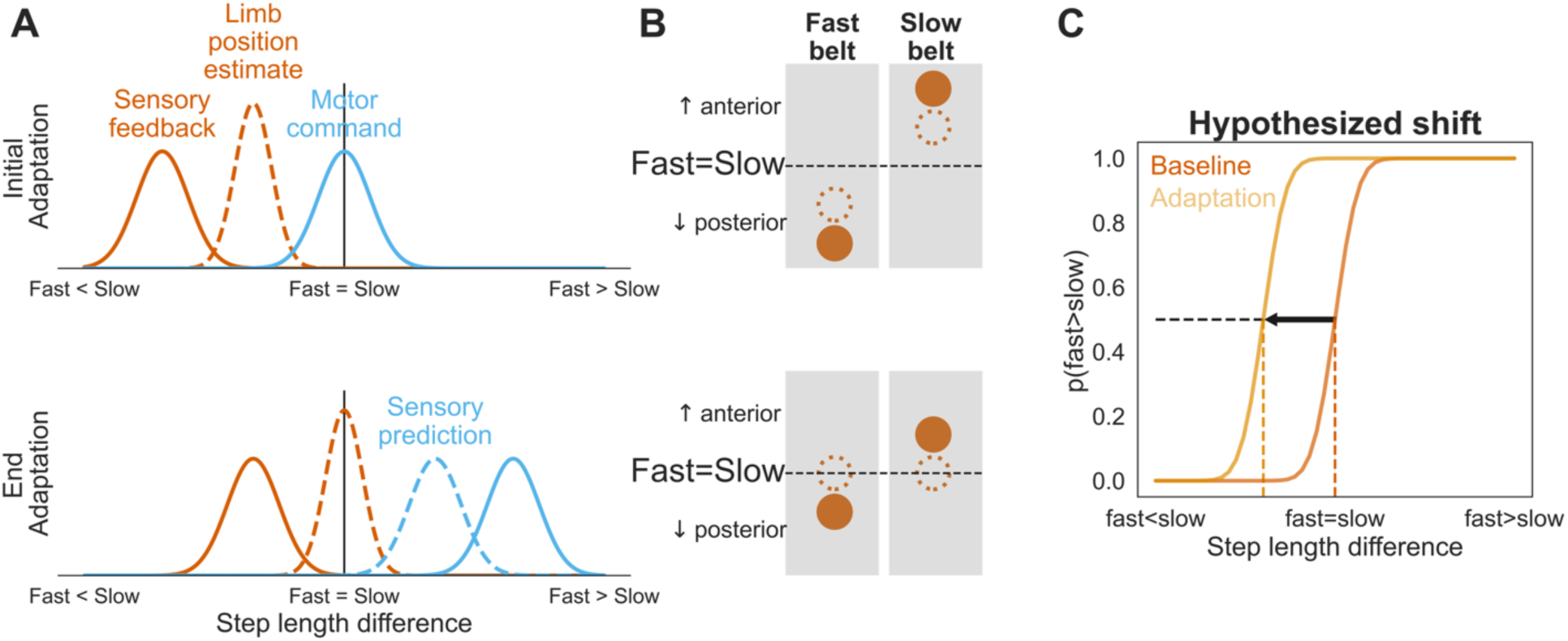
Theory and hypothesis. **(A)** Optimal integration of the motor command and sensory feedback. At initial adaptation (top) the sensory prediction of the motor command (hidden under the solid blue function) and sensory feedback of the limb positions (orange function) are misaligned. The resulting limb position sense estimate (dashed orange function) would be biased towards the sensory prediction. Adaptation would eventually reach a plateau (bottom) when the position sense estimates of the fast and slow limbs are equal. **(B)** Depiction of how the proposed sensory recalibration in (A) would be represented in terms of each individual limb when viewed from above. The fast limb should feel more anterior and/or the slow limb should feel more posterior compared to their true positions. **(C)** Hypothetical depiction of how the shift in (A) and (B) would be represented in the context of the position sense test we used in the current experiment. Participants performed a 2- alternative forced choice (2-AFC) task making decisions between either the fast or slow limb being more anterior. Therefore, this task provided a measure of where the fast and slow limbs are perceived as equal (i.e., the position of subjective equality, or PSE). According to our hypothesis, after adaptation, the PSE should shift to the left, indicating that while the fast limb is actually behind the slow limb, they are reporting it being further ahead relative to baseline.

Based on this framework, we generated 3 hypotheses: First, limb position sense should shift such that the fast limb feels more anterior and/or the slow limb feels more posterior than the true position (Figure 1C). Second, the magnitude of this sensory recalibration should be related to the plateau of adaptation, such that the greater the sensory recalibration the less the adaptation plateau. Third, the uncertainty inherent in the sensory feedback at baseline, should be related to the adaptation plateau. This is because we assume the sensory prediction and sensory feedback of the limb position are optimally combined to form the limb position estimate. Thus, if there is greater sensory uncertainty, there will be a greater sensory recalibration towards the sensory prediction, resulting in a lower adaptation plateau.

We tested these hypotheses by measuring lower limb position sense with a previously validated position sense task [23] before and after a long bout of split-belt walking (∼45 minutes). The data did not support the proposed theoretical model of sensory recalibration during split-belt adaptation: We did not observe sensory recalibration and neither the individual sensory shifts nor the baseline sensory uncertainty were related to the plateau of adaptation as measured by step length asymmetry. However, post-hoc analysis of other measures of split-belt adaptation, specifically of double limb support time, revealed a strong relationship between its adaptation plateau and sensory recalibration. These results suggest that that sensory recalibration could play an important role in split-belt adaptation and raise questions regarding the error signal driving split-belt adaptation.

## Materials and Methods

Here, we tested sensory recalibration (a shift in lower limb position sense estimates) during split-belt adaptation. We used a standard split-belt paradigm to measure sensorimotor adaptation during gait [1,24]. We measured lower limb position sense estimates using a previously validated and reliable 2-alternative forced choice (2-AFC) task [23].

### Participants

Seventeen young healthy adults between the ages of 18-35 years, (5 female) participated in the study (recruited between 11/11/2022 to 5/16/2023). Participants were excluded if they reported any current or recent musculoskeletal or neurologic diagnoses, dizziness, pain, or anything that might alter either sensation in the lower limbs or motor learning. All participants gave their written informed consent before participating in the study. This study was reviewed and approved by the University of Delaware Institutional Review Board.

### Experimental procedure

Participants completed a single study session comprising 3 walking phases and 3 position sense tests. Throughout the testing, participants wore a ceiling-mounted safety harness that did not provide body-weight support and held on to the safety bar in front of them throughout the study (Figure 2A). Participants wore a black drape throughout the task, which obstructed vision of the lower limbs. During the position sense tests, they also donned a pair of noise-reducing headphones to mitigate the influence of auditory feedback on their decisions.

**Figure 2.**
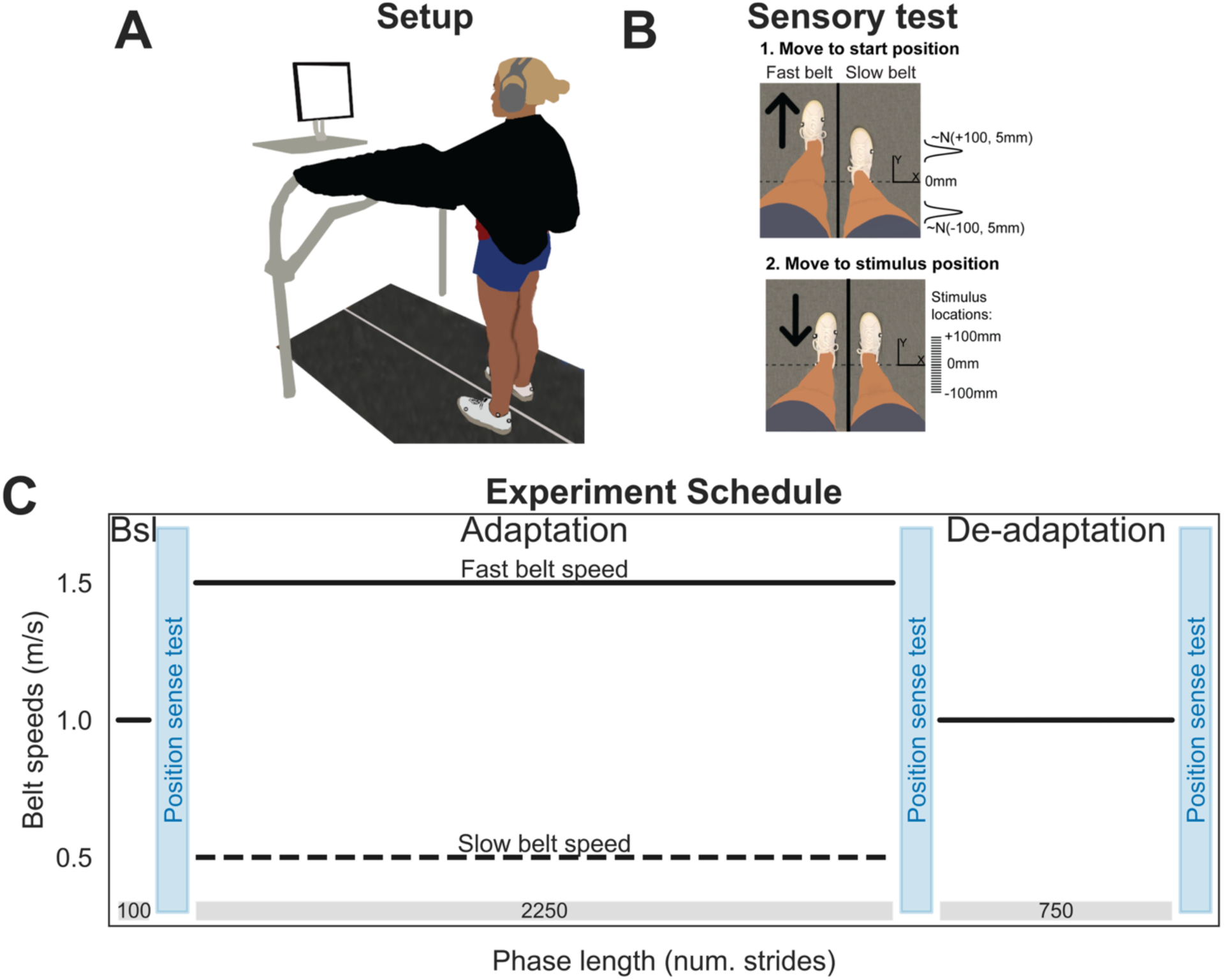
Experimental paradigm. **(A)** Task setup. Both the split-belt adaptation task and position sense tests were performed on a split-belt treadmill without vision of their feet. The computer monitor provided task feedback only during the position sense tests. **(B)** Depiction of a single trial for the position sense test trial. First, the fast limb was moved by the treadmill belt to a start position, randomized to be either in front or behind the slow limb (which remained stationary throughout the test). The exact start position of the fast limb was sampled from a normal distribution centered on either 100mm ahead or behind of the slow limb. Second, the fast limb was moved by the treadmill to one of 21 different stimulus locations selected based on an adaptive Bayesian algorithm (see Methods). **(C)** Experimental schedule. Participants performed baseline (Bsl), adaptation and de-adaptation phases in sequential order. Position sense tests (blue bars) were performed after each phase. During the adaptation phase, the fast belt moved at 3x the speed of the slow belt. The length of each phase (num. strides) is denoted in the grey box. **A** and **B** were adapted from [23] under the terms of the CC BY 4.0 license (http://creativecommons.org/licenses/by/4.0/).

We used a standard split-belt adaptation paradigm, consisting of baseline, adaptation, and de-adaptation phases (Figure 2C). During the baseline phase (100 strides, ∼2 minutes) both belts moved at 1.0 meters / second. Immediately after the baseline phase, participants performed the first position sense test which served as their baseline measure of limb position. Next, participants performed the adaptation phase (2250 strides, ∼45 minutes) where the left belt moved at 1.5 m/s and the right belt moved at 0.5 m/s, a 3:1 ratio. We performed a long adaptation phase so that participants could achieve a stable plateau of their adaptation behavior [24]. Immediately after the adaptation phase, participants performed the second position sense test. Next, participants performed a de-adaptation phase (750 strides, ∼15 minutes) where both belts returned to moving at 1.0 m/s. Immediately after the de-adaptation phase, participants performed the third position sense test to determine if their perception of limb symmetry returned to baseline or was maintained after de-adaptation [11,12].

We assessed lower limb position sense using a 2-AFC task [23], where participants were asked to report if they felt their left or right foot was more forward (i.e., anterior), after the fast limb was passively moved by the treadmill to one of 21 possible stimulus positions. Stimulus positions were measured as the difference between the fast and slow limb’s ankle markers in the y-axis of the lab, in millimeters. Positive values indicate the fast limb was more forward while negative values indicate the slow limb was more forward. This allowed us to determine the specific position at which participants perceived their feet to be in equal position relative to the lab’s y-axis. Only the limb on the fast belt was moved throughout the task, while the limb on the slow belt remained stationary. Each position sense test consisted of 50 trials with a seated rest break provided after the 25^th^ trial. After the break, the experimenter manually placed their feet back in the same position as before the break. Participants were asked not to move their limbs or shift their body weight during this task, and this instruction was monitored visually by the experimenter using a custom GUI which received data from the force plates.

Each trial in the position sense test consisted of two steps (Figure 2B): 1) a movement to a start position (at a uniformly random speed between 40 and 50 mm/s) 2) a movement to a stimulus position (at a uniformly random speed between 10 and 30 mm/s). The start position was used to prevent the location of prior stimulus positions from influencing decisions. We ensured the limb moved to the stimulus position both forward and backward an equal number of times to prevent a bias in the responses due to movement direction [23]. The start positions themselves were sampled from a normal distribution with a mean of either -100 mm or +100 mm of foot position difference, depending on the movement direction for that trial, and a standard deviation of 5 mm. The treadmill paused at the start position for a uniformly random time between 0-2 seconds before moving to the stimulus position. The stimulus positions were selected from 21 possible foot position differences: every 10 mm between -100 to +100 mm. The majority of stimuli were selected by an adaptive algorithm (see below). Once the fast limb reached the stimulus position, participants were asked “Do you feel your right or left foot is more forward”. Responses were recorded by the experimenter.

### Adaptive algorithm

We used a Bayesian adaptive algorithm to efficiently estimate both the lower limb position sense and the sensory uncertainty. We describe only the essential concepts of the adaptive algorithm here; further details including equations can be viewed in our prior paper [23]. First, we assumed that the probability of responding that the fast limb was more forward for each participant took the form of a psychometric function, specifically, a cumulative normal distribution with the mean (μ) and standard deviation (σ). Therefore, μ represents the point of subjective equality (PSE) or the location where participants judge their feet as equal along the laboratory’s y-axis, and σ reflects the uncertainty within the proprioceptive system. Both these parameters were estimated on each trial by the adaptive algorithm using Bayes rule [38]. All possible values of μ and σ represent a joint probability distribution that is updated after the response on each trial. This distribution is estimated by multiplying the Prior (i.e., the probability of all possible μ and σ values) with the Likelihood (the probability of responding “left” or “right” given all possible μ and σ values). The resulting Posterior becomes the Prior for each successive trial. We initialized the prior for μ as a 0-mean normal distribution with a standard deviation of 20, and for σ as an exponential distribution with an expected value of 20 [23]. The Likelihood is represented as the probability of the participants actual response, given all possible values of μ and σ, across all stimulus positions. We normalized the Posterior, then marginalized across the two dimensions. The mean of each marginalized distribution served as our estimate of μ and σ parameters on each trial. The parameter estimates after trial 50 were our final μ and σ estimates for that position sense test, representing the PSE and sensory uncertainty, respectively.

We also used the adaptive algorithm to select the most informative next stimulus position to improve the efficiency of the task [38]. This was accomplished this using the principle of information entropy, a measure of uncertainty in a given distribution [39]. Before each trial, we calculated the expected reduction in uncertainty in the Posterior distribution at each possible stimulus position. The stimulus position that reduces uncertainty the most is definitionally the stimulus position that will increase the amount of information gained on that trial and thus the most efficient stimulus position to probe on the next trial. We simulated the posterior probability distribution for both possible responses (i.e., left and right) on the next trial, at every possible stimulus position. The combined expected reduction in entropy is then calculated as a weighted average of expected entropy for each response (left and right). The stimulus position that minimizes the expected reduction in entropy was selected on each trial.

While these stimulus positions are highly efficient, in practice participants can become mentally fatigued by the difficulty of constantly judging between stimuli this close together. Therefore, to keep participants engaged in the task, we injected stimuli that were less challenging (except for the first 5 trials). These pre-selected stimuli were either far stimuli (± 100 or ± 90), injected randomly once every 10 trials, or near stimuli (±10, ± 20 or ± 30 mm from the current PSE estimate, rounded to the nearest stimulus location), injected randomly once every 5 trials. This modified version of an adaptive Bayesian algorithm can reliably estimate both parameters in as little as 50 trials [23].

### Data collection

A split-belt treadmill with dual motors and separate force plates was used throughout this experiment (Bertec, Columbus, OH). Kinematic data were collected using eight Vicon MX40 cameras and Vicon Nexus software (version 2.8.2, London, England). We used a custom marker set of seven retroreflective markers, located on the participant’s heels, 5th metatarsals, and lateral malleoli on both feet. An additional marker was placed on the subject’s left 1st metatarsal so the system could determine the difference between the left and right feet during data collection. Commands for the split-belt treadmill during the walking task and the position sense test were controlled using custom-written Matlab software (version 2022a, MathWorks, Natick, MA). The minimal data set and accompanying code, where generated is available at The Open Science Framework via https://doi.org/10.17605/OSF.IO/VQBFH

### Data analysis

For the kinematic data during the walking phases, gaps in marker trajectories were filled using Woltring filter for small gaps (one to four frames) and Pattern Fill for larger gaps (more than four frames) in Nexus. Next, custom-written Matlab scripts were used to filter the kinematic data and select gait events. The kinematic data were low pass filtered at 10Hz using a fourth order Butterworth filter. Heel strike was determined as the frame when the heel marker velocity transitioned from positive to negative, and toe off events were determined as the frame when the heel marker velocity transitioned from negative to positive [40]. These events were used to calculate step length and double limb support time.

Left and right step lengths were calculated as the difference between the ankle markers at each limb’s respective heel strike, along the y-axis of the lab. Any step length 3x larger than the inter-quartile range for that participant was removed from the remainder of the analysis. Step length was then used to calculate step length asymmetry on each stride, the difference between the fast and the slow limb, normalized by their sum. We express this measure as a percentage where 0% indicates the limbs are perfectly symmetrical, positive values indicate the fast limb is stepping longer than the slow limb and negative values indicate the fast limb is stepping shorter than the slow limb. Step length asymmetry was baseline corrected for each participant by subtracting the mean of the final 50 strides of the baseline phase from all strides of the experiment. The adaptation plateau was calculated as the mean step length asymmetry of the final 50 strides of adaptation. We hypothesized that the adaptation plateau measure should be related to sensory recalibration and baseline sensory uncertainty (see below).

Based on our main results, we performed a series of post-hoc analyses using different measures of split-belt adaptation: the baseline corrected fast and slow step length and baseline corrected double limb support time. Double limb support time for each limb was calculated as a percentage of each stride cycle. Specifically, the fast double limb support time was calculated as the time between the slow limb heel strike and the fast limb toe off, divided by the total time of that stride and expressed as a percentage. The slow double limb support time was calculated as the time between the fast limb heel strike and slow limb toe off, divided by the total time for that stride and expressed as a percentage. To characterize the adaptation plateau for these measures we again used the mean of the final 50 strides of the adaptation phase.

For our measures of lower limb position sense, we used the final μ and σ estimates from the adaptive algorithm (i.e., after trial 50). The estimate of μ represents the location in which the participant perceives their feet as being symmetrical along the lab’s y-axis, or PSE. We characterized sensory recalibration by subtracting the PSE estimates measured after baseline from those measured after adaptation. We also performed another position sense test after the de-adaptation phase to determine if any shift in the limb position changed after de-adaptation compared to after adaptation (PSE after de-adaptation minus PSE after adaptation) [11,12]. The estimate of σ represents the uncertainty or noise in the sensory system. We characterized baseline sensory uncertainty as the σ value from the position sense test measured after the baseline phase.

### Statistical analysis

Statistical analyses were both performed using custom written Python notebooks (v4.3.0). We used a Bayesian approach to make statistical inferences [41], estimating the data generating process using two different statistical models. For the hypothesis that sensory recalibration occurs as a result of split-belt adaptation, we assumed that each individual PSE estimate (i) was generated from a normal distribution with a mean depending on experimental phase (α_phase_; post-baseline, post-adaptation and post-de-adaptation) and subject (α_subj_):

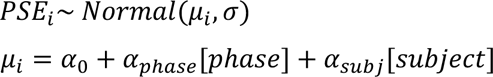

An additional Bayesian regression model was used to test our other hypotheses regarding the relationships between variables:

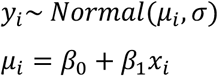

Here, we again assume that each participant’s (i) predictor variable (y, the adaptation plateau) is sampled from a normal distribution with a μ_i_ that depends on an intercept (β_0_) and slope (β_1_). And, x is the predictor variable (sensory recalibration or uncertainty).

For both models, we estimated the posterior distributions for each parameter in this model using Bayes rule, combining evidence from our data with our prior assumptions regarding each parameter value. The priors were set conservatively so that they did not unduly influence the posterior [41]. We estimated the joint posterior distribution using the PyMc 4 library [42] and the bayes-toolbox Python package [43]. We used Markov Chain Monte Carlo sampling to sample from joint posterior distribution 10,000 times with 2,000 tuning samples across 4 chains. We performed posterior predictive checks to ensure that the posterior samples accurately represented the data [41,44].

We report the full range of credible effects, focusing on the difference between the phase variable for the first model (α_phase_ parameters) and the beta coefficient for slope (β_1_) for the second model. To make inferences we used the 95% high density interval (HDI) defined as the narrowest span of credible values that contain 95% of the distribution [41] as well as the probability of a difference, defined as the percent of the posterior parameter distribution on one side of 0. The HDI reflects the size of the effect with 95% certainty while the probability of a difference reflects the certainty of the effect, with larger values being more certain. We do not provide a specific decision threshold for the latter measure as we would with an alpha value in frequentist statistics, allowing for a more nuanced interpretation of the results.

## Results

Seventeen (5 female) participants (mean age [95% HDI] = 23.9 years [21.1 26.6]) completed the experiment. Figure 3A displays the group averaged step length asymmetry timeseries data. Typical of split-belt adaptation paradigms, the split-belt initially produced a large negative step length asymmetry, indicating the fast limb was stepping shorter than the slow limb. As the adaptation phase continued, the step lengths became more symmetrical but then adapted past baseline symmetry, denoted by the positive step length asymmetry index at the end of baseline. This finding is similar to a prior study with an adaptation phase of this length and is hypothesized to occur because it is more energetically optimal [24]. During the de-adaptation phase, when the belts returned to tied, participants displayed the typical aftereffect in the opposite direction of the initial perturbation. The robust aftereffect indicates the novel stepping pattern was stored (i.e., the forward model was recalibrated), despite the additional time required between the phases to perform the position sense test.

**Figure 3.**
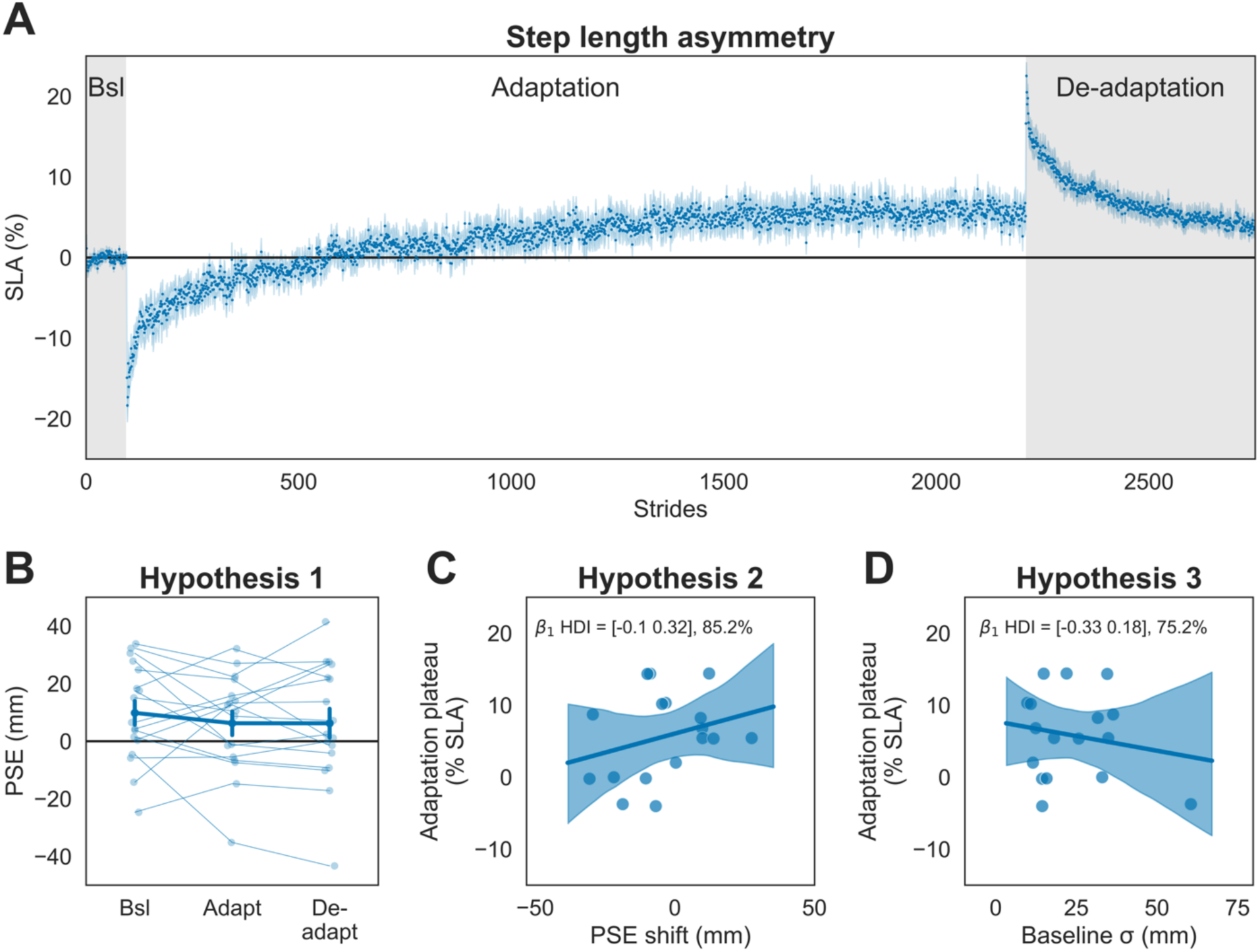
Split-belt adaptation does not induce sensory recalibration. **(A)** Group averaged step length asymmetry data across the experiment (Bsl = Baseline). Each dot represented the group-averaged step length asymmetry for one stride, and the shading represents 1 standard error of the mean. **(B)** Hypothesis 1. We did not observe sensory recalibration at the group level as the estimated PSEs for each position sense test did not shift across the three phases. The data for each participant across the tests are represented as the thin, connected dots. The overall mean is represented as the thick line. Error bars represent 1 standard error of the mean. **(C & D)** Hypothesis 2 & 3. Sensory recalibration was not related to the adaptation plateau for step length asymmetry, nor was the baseline sensory uncertainty. Each plot represents one participant as a dot. The regression lines were created with the mean Bayesian regression parameters, and the shading represents the 95% HDI. The HDI and p_difference_ values for the slope parameter, β_1_, are provided above the plots.

Our first hypothesis was that limb position sense estimates would shift in the negative direction from baseline to adaptation. Figure 3B displays participant and group averaged PSE estimates across the three timepoints. We did not observe a reliable shift in PSE estimates from baseline to adaptation (mean [95% HDI] PSE difference = -0.46 mm [-2.54 1.32], p_difference_ = 75.1%). Additionally, there was no shift in the PSE from the adaptation to de-adaptation phase (- 0.003 mm [-1.86 1.89], p_difference_ = 50.0%). This lack of a difference was not due to changes in uncertainty after adaptation from baseline (0.26 mm [-1.71 2.34], p_difference_ = 63.8%), which is consistent with a prior split-belt adaptation study [18]. Therefore, the hypothesis that limb position sense estimates would shift during split-belt adaptation was not supported.

While there was no group level shift in limb position sense estimates, sensory recalibration could still predict the adaptation plateau at the participant level (Figure 3C). We hypothesized there would be a positive relationship between sensory recalibration and the adaptation plateau for step length asymmetry. While this relationship was in the predicted direction, it was not reliable (β_1_ mean [95% HDI] = 0.11 [-0.10 0.32], p_difference_ = 85.2%). Similarly, we did not find support for the hypothesis that baseline sensory uncertainty was related to the adaptation plateau (Figure 3D; β_1_ = -0.08 [-0.33 0.18], p_difference_ = 75.2%). Combined, these results do not support the theory of optimal integration between the sensory feedback and sensory prediction driving sensory recalibration during split-belt adaptation. Indeed, the finding that step length asymmetry adapted beyond baseline asymmetry (plateau > 0) is inconsistent with the overall theoretical framework we proposed.

### Post-hoc analyses

Despite the lack of support for the optimal integration theory we proposed, it still is possible that shifting position sense estimates play an important role in split-belt adaptation. These analyses were motivated by both the large amount of variability we observed in the PSEs from baseline to adaptation (Figure 3B), and the fact that the primary error signal driving split-belt adaptation is not known as multiple measures robustly adapt to this perturbation [1,14,15,24,45]. Because of our limited marker set – which we selected to prevent additional sensory feedback during the position sense tests – we were able to assess individual step lengths and double limb support times. We performed post-hoc analyses to assess relationships between sensory recalibration and the adaptation plateau measured with these gait parameters.

Figure 4A displays the group averaged timeseries data for baseline corrected step lengths. Here, we observe that the fast limb’s step length returns to baseline by the end of adaptation. Conversely, the slow limb’s step length quickly adapts past baseline and remains there, plateauing at a range well below zero. Thus, the fact that step length asymmetry adapted well beyond baseline (Figure 3A), can be attributed to the slow limb rather than the fast limb. In Figure 4C and D, we display the relationship between sensory recalibration and the adaptation plateau for each limb’s step length. We found a reliable relationship between sensory recalibration and the adaptation plateau for the fast step length (β_1_ = 1.16 [-0.14 2.48], p_difference_ = 95.9%), but not for the slow step length (β_1_ = 0.04 [-1.32 1.42], p_difference_ = 51.5%). Like the step length asymmetry analysis, baseline sensory uncertainty could not reliably predict the adaptation plateau for either step length (fast β_1_ = -0.36 [-2.27 1.39], p_difference_ = 66.1%; slow β_1_ = 0.23 [-1.58 1.95], p_difference_ = 60.9%).

**Figure 4.**
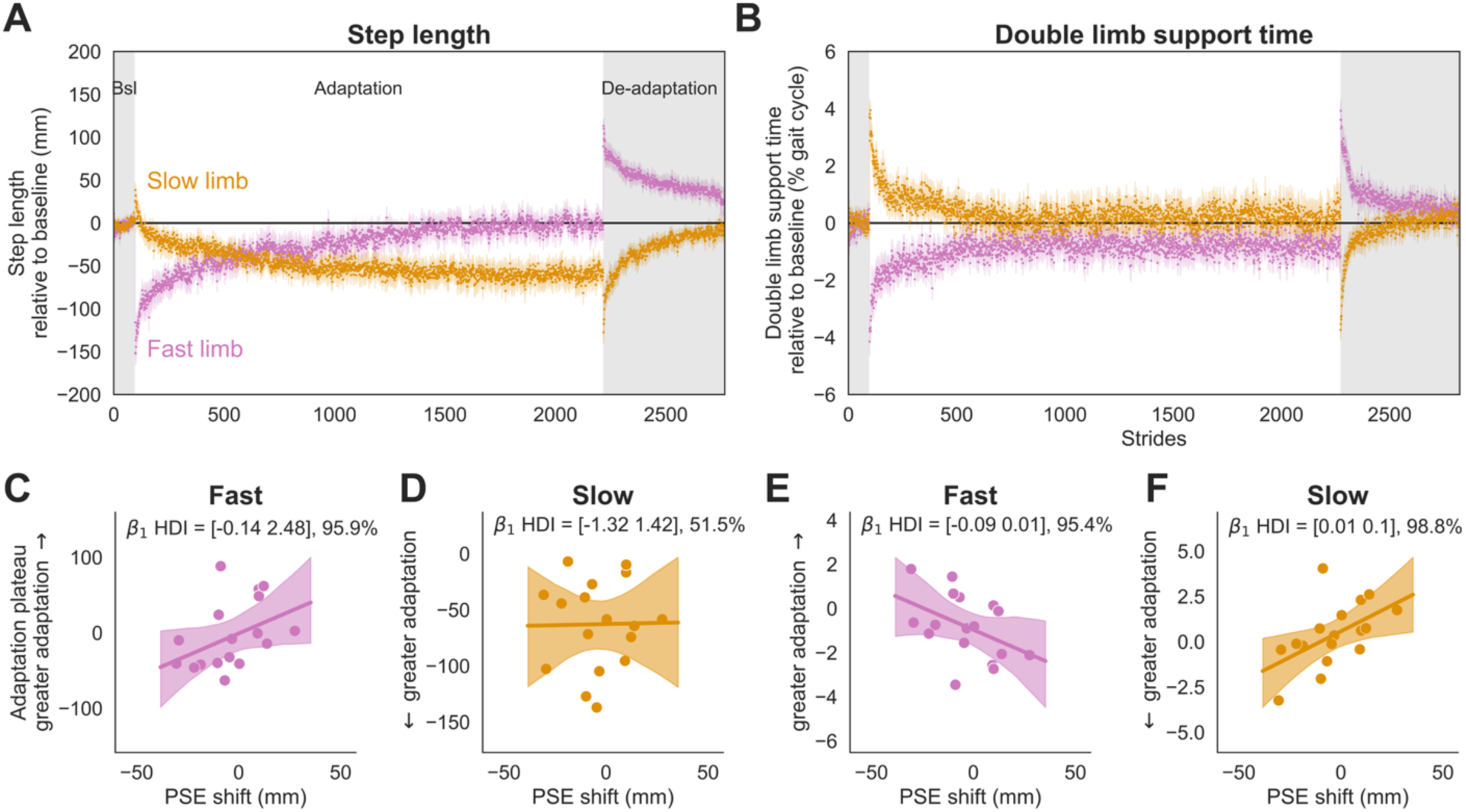
Fast and slow step length and double limb support time adaptation data. **(A&B)** Group averaged fast and slow step length and double limb support time (Bsl = Baseline). Each dot represented the group-averaged metric for one stride, and the shading represents 1 standard error of the mean. Note that each of these measures is baseline corrected **(C-F)** Relationships between sensory recalibration and the adaptation plateau for each gait parameter. Each plot represents one participant as a dot. The regression lines were created with the mean Bayesian regression parameters, and the shading represents the 95% HDI. The HDI and p_difference_ values for the slope parameter, β_1_, are provided above the plots. **(C)** Fast step length. **(D)** Slow step length. **(E)** Fast double limb support time **(F)** Slow double limb support time.

We also assessed double limb support time. The average baseline corrected double limb support time for each limb is plotted in Figure 4B. Interestingly, neither the fast nor the slow double limb support time adapted past baseline, despite the extended period of adaptation. Additionally, sensory recalibration was reliably related with the adaptation plateaus for both the fast (β_1_ = -0.04 [-0.09 0.01], p_difference_ = 95.4%) and slow (β_1_ = 0.06 [0.01 0.11], p_difference_ = 98.8%) double limb support times (Figure 4 E&F). Again, baseline sensory uncertainty was not related to adaptation plateau for double limb support time for either limb (fast β_1_ = -0.01 [-0.08 0.05], p_difference_ = 66.2%; slow β_1_ = -0.02 [-0.09 0.04], p_difference_ = 76.8%). Therefore, when split-belt adaptation is characterized in terms of double limb support time, sensory recalibration is consistently related to the adaptation plateau.

## Discussion

The purpose of this study was to assess sensory recalibration during split-belt adaptation. Guided by a theoretical model of optimal integration between sensory feedback and the sensory prediction, we tested three hypotheses. First, lower limb position sense would shift after adaptation relative to baseline (i.e., sensory recalibration). Second, sensory recalibration would be related to the adaptation plateau. And third, baseline sensory uncertainty would be related to the adaptation plateau. None of these hypotheses were supported by the data when split-belt adaptation was assessed as step length asymmetry. However, our post-hoc analyses revealed a relationship between sensory recalibration and the adaptation plateau when measured as the fast limb step length or fast and slow double limb support time. This finding indicates that sensory recalibration is involved in split-belt adaptation, and the double limb support time data in particular raise questions about the primary error signal that is driving adaptation.

We reasoned that sensory recalibration could occur during split-belt adaptation by optimally integrating sensory feedback of limb positions and the predicted sensory consequences of the motor command (i.e., the sensory prediction) which should be misaligned at the start of split-belt adaptation. We based our hypothesis on studies suggesting that an efference copy is used to generate sensory predictions of the motor command, and this sensory prediction is optimally combined with the sensory feedback from the periphery to form a unified percept of the limb locations [22,46,47]. Prior work has incorporated this theory into computational models of sensorimotor adaptation during reaching [27,31–33]. These models were designed for visuomotor adaptation paradigms where the mismatch between visual and proprioceptive feedback is thought to be the primary driver of sensory recalibration. Importantly, the difference between visual and proprioceptive feedback is maintained throughout this adaptation paradigm. For split-belt adaptation, the difference between sensory feedback and the sensory prediction is likely not consistent throughout the adaptation phase. This is because the forward model – the expected sensory consequences of the motor command – is recalibrated. Thus, one possible explanation for the lack of sensory shift in the current study is that the discrepancy driving sensory recalibration could be resolved prior to the adaptation plateau, a possibility that still fits under the optimal integration framework described in the current study.

Of course, it is also possible that part or all the optimal integration framework we proposed is incorrect. One key assumption of our theoretical model was that the error signal being corrected was the perceived difference between the left and right limb positions, similar to the proprioceptive error hypothesis [27]. This contradicts previous theories of sensorimotor adaptation which hypothesize that the difference between the sensory prediction and the sensory feedback drives adaptation (i.e., the sensory prediction error hypothesis) [36]. Optimal integration driving sensory recalibration as we hypothesized in the current study is incongruent with the sensory prediction error hypothesis. This is because, at the adaptation plateau, the difference between the sensory prediction and sensory feedback difference would be fully corrected. Continuing with this line of reasoning, we would not expect to observe a relationship between baseline sensory uncertainty and the adaptation plateau or between the sensory recalibration and the adaptation plateau. While we observed the latter relationship when measuring adaptation as double limb support time, we did not observe the former relationship for any measure of adaptation. Taken together, we think the most parsimonious explanation of the current findings is that optimal integration of the sensory prediction and sensory feedback does not drive sensory recalibration during split-belt adaptation.

Another explanation of the current findings is that sensory recalibration is not involved in split-belt adaptation [25], a theory based on prior studies that did not observe reliable lower limb position sense shifts [18,19]. Despite using a validated version of the lower limb position sense estimates and assessing these estimates after a long bout of split belt adaptation, the current study also did not observe sensory recalibration at the group level. However, this prior work did not assess the relationship between sensory recalibration and the adaptation plateau, which when measured as double limb support time, provides evidence against the theory that sensory recalibration is not involved in split-belt adaptation. Indeed, this observation suggests that sensory and motor recalibration are not independent processes.

Therefore, one possible alternative to the theory that sensory recalibration is not involved in split-belt adaptation, and to the optimal integration framework we proposed here is that sensorimotor adaptation drives changes in the somatosensory system to reduce the felt perturbation [48]. The relationship between double limb support time and sensory recalibration supports this hypothesis: Early in split-belt adaptation, the fast double limb support time – which essentially reflects how long the fast limb stays on the treadmill relative to the slow limb’s heel strike – is much shorter compared to baseline. This is expected as the fast belt is quickly forcing the limb further into extension during the stance phase [14]. Therefore, the position of the limb at toe off and double support time are closely linked, making it possible for the perceived limb position to be related to double limb support time. To help resolve this perturbation, the fast limb could be perceived as more anterior relative to baseline. Thus, the greater the adaptation plateau of double limb support time the more anterior the fast limb might be perceived (a negative shift).

Interestingly, we did not observe a relationship between sensory shift and adaptation plateau for the typical measure of split-belt adaptation, step-length asymmetry. We selected this measure because it is ostensibly related to limb position sense and it is frequently used as a measure of split-belt adaptation. However, the error signal driving split-belt adaptation is not known; rather, several inter-related features of gait demonstrate adaptation-like behavior during and after split-belt treadmill walking (e.g., step length, double limb support time, center of pressure, and at a more granular level, muscle firing patterns) [1,14,15,49]. Step length asymmetry has received perhaps the most attention, but given enough time this gait measure consistently adapts past baseline symmetry (i.e., the presumed “goal”) [24], a finding that was replicated in the current study. Adapting past the goal has not been observed in any other form of sensorimotor adaptation to our knowledge and is not predicted by any computational models of sensorimotor adaptation [27,31,50,51]. Given this over-adaptation of step length asymmetry, Sánchez and colleagues hypothesized that energy optimization is the signal driving split-belt adaptation, specifically of the fast limb using the positive work from the treadmill [24,45,52]. However, if the fast limb is driving the over-adaptation, we would have expected to observe that it adapted past baseline rather than the slow limb. Instead, we observed the slow limb quickly adapted past baseline early in the adaptation phase and maintained that pattern until the end of adaptation.

We propose that double limb support time is a metric that is more closely related to the true error signal of split-belt adaptation. The first reason for this claim is that we did not observe over-adaptation past baseline for either fast or slow double limb support times, even after a very long adaptation phase. This is consistent with measures of sensorimotor adaptation in other paradigms [53–59]. Second, the adaptation plateau of double limb support time was the only parameter that was reliably related to the sensory shift, and sensory shifts and adaptation plateaus are related in other sensorimotor adaptation paradigms [27,28,60]. Third, double limb support time has face validity as a critical objective for adaptation because it is an important measure for stability of gait. It specifically reflects the timing of the shift to support the body weight from one limb to the other. For example, fast double limb support time reflects the percent of the gait cycle that it takes to accept the weight of the body, as the support shifts from the fast limb to the slow, a key measure of stable gait. Interestingly, another gait measure that is closely related to stable gait, center of pressure asymmetry, also robustly adapts during split-belt adaptation [15]. We speculate that double limb support time reflects some critical measure of stability that is the primary error signal driving split-belt adaptation.

Overall, the hypothesized framework for sensory recalibration during split-belt adaptation – optimal integration of sensory feedback and the sensory prediction driving sensory recalibration during split-belt adaptation – was not supported by the data. Still, we contend that sensory recalibration is involved in split-belt adaptation based on our post hoc analyses of double limb support time. While the specific nature of this relationship is unclear, limb position sense should be considered in models of split-belt adaptation given the importance of proprioception to stable gait.

